# Gene expression profiles and pathway enrichment analysis to identification of differentially expressed gene and signaling pathways in epithelial ovarian cancer based on high-throughput RNA-seq data

**DOI:** 10.1101/566331

**Authors:** A Siavoshi, M Taghizadeh, E Dookhe, M Piran

## Abstract

Epithelial ovarian cancer (EOC) can be considered as a stressful and challenging disease among all women in the world, which has been associated with a poor prognosis and its molecular pathogenesis has remained unclear. In recent years, RNA Sequencing (RNA-seq) has become a functional and amazing technology for profiling gene expression. In the present study, RNA-seq raw data from Sequence Read Archive (SRA) of six tumor and normal ovarian sample was extracted, and then analysis and statistical interpretation was done with Linux and R Packages from the open-source Bioconductor. Gene Ontology (GO) term enrichment and Kyoto Encyclopedia of Genes and Genomes (KEGG) pathway analysis were applied for the identification of key genes and pathways involved in EOC. We identified 1091 Differential Expression Genes (DEGs) which have been reported in various studies of ovarian cancer as well as other types of cancer. Among them, 333 genes were up-regulated and 273 genes were down-regulated. In addition, Differentially Expressed Genes (DEGs) including RPL41, ALDH3A2, ERBB2, MIEN1, RBM25, ATF4, UPF2, DDIT3, HOXB8 and IL17D as well as Ribosome and Glycolysis/Gluconeogenesis pathway have had the potentiality to be used as targets for EOC diagnosis and treatment. In this study, unlike that of any other studies on various cancers, ALDH3A2 was most down-regulated gene in most KEGG pathways, and ATF4 was most up-regulated gene in leucine zipper domain binding term. In the other hand, RPL41 as a regulatory of cellular ATF4 level was up-regulated in many term and pathways and augmentation of ATF4 could justify the increase of RPL41 in the EOC. Pivotal pathways and significant genes, which were identified in the present study, can be used for adaptation of different EOC study. However, further molecular biological experiments and computational processes are required to confirm the function of the identified genes associated with EOC.

## 1. Introduction

Today, Epithelial Ovarian Cancer (EOC) [1] can be considered as a stressful and challenging disease among all women in the world. EOC is diagnosed annually in nearly a quarter of a million women, and it is responsible for 140,000 deaths each year [2]. Some of the most significant causes of more than a one-half of patient death are the lack of diagnosis and validated clinical applicable test to effective therapies in the early stages, different types of ovarian cancer in the female reproductive system[3], [4], [5] and also resistant to chemotherapy of the disease [6], [7]. At present, the commonly used treatment approaches of EOC include surgical resection, radiotherapy, chemotherapy, blood tests for investigation of biomarkers such as CA125, HE4 that improve patients’ prognosis and recurrence[8]. Nevertheless, survival rate of EOC due to Sensitivity and specificity of histological criteria is still low, especially in advanced EOC [9]. The major research project goal is the investigation and discover DEGs related to a specific biological function between normal and tumor ovarian samples. Until now, there has not been any remarkable success to treatment of ovarian cancer, because of the lack of clear understanding of the changes in the molecular variability of ovarian cancer at different levels of the disease. Recently, some researches on finding target genes, in order to introduce as a biomarker for early detection of ovarian cancers, have been impressive [10], [11], [12]. However, recent transcriptome analysis on RNA-Seq [13] data have shown great promise for the discovery of new and more useful biomarkers [14], [15]. RNA-Seq is a powerful genomic tool that can explain the relationship between clinical features and their biological changes for new treating strategies of ovarian cancer, and describe the mechanism of the disease according to the gene expression process at each stage and level of the disease [16], [17], [18].

RNA-Seq is generally used to compare gene expression between conditions, such as drug treatment versus non-treated, and find out which genes are up- or down-regulated in each condition. Many packages are available for this type of analysis, some of the most commonly used tools are DESeq2 [19] and EdgeR [20] packages from Bioconductor (http://www.bioconductor.org/) [21]. Both of these tools use a model based on the negative binomial distribution [22]. Various studies have shown that comparing the level of gene expression among thousands of genes between normal and cancerous samples. RNA-Seq is a powerful tool to investigation of the cancer pathogenesis and development of new drug agents to cancer treatment [23], [24], [25]. Differential Expression (DE) [26], [27] analysis as the most important applications of RNA-Seq data analysis can be used for downstream systems biology analysis such as gene ontology (GO) [28], [29] to provide insights into cellular processes altered between biological function and Enrichment of biological pathways supplied by Kyoto Encyclopedia of Genes and Genomes (KEGG) [30]. These analyses are routinely used in a variety of research areas including type of cancer and human disease research such as ovarian cancer [31], lung, prostate, bladder, colorectal, gastric [32], [33], [34], [35], [36] and drug development [37].

The purpose of the present study was to compared the ovarian cancer RNA-seq data in two levels of normal and tumor ovarian for the presentation of different genes expressed by the use of statistical analysis and performing enrichment pathways by using GO and KEGG analysis. To develop new biomarkers through such analyses, identification of key differential gene expression (DEG) for diagnosis and prognosis in each level of EOC will be very useful.

## 2. Materials & Methods

### 2.1. Data processing method and software sources

RNA-Seq raw data (SRR3289902), Cristofer, et al [38] was downloaded from the Sequence Read Archive (SRA) (www.ncbi.nlm.nih.gov/geo) in format of Fstaq. The data included three ovarian tumor and three normal samples. Ubuntu 17.10 (64-bit) was used to process raw data and Statistical software R (version 3.5.1, https://www.r-project.org/) was used to statistical calculation and interpretation of DEGs. Biological significance of DEGs was explored by GO term enrichment analysis including biological process (BP), cellular component (CC), and molecular function (MF), based on Bioconductor packages enrichR (https://cran.r-project.org/package=enrichR) and then KEGG pathway enrichment analysis of DEGs was performed with enrichR as well.

### 2.2. RNA sequencing data Analysis

#### 2.2.1. Quality control, read mapping and Counting reads in features

Quality control and preprocessing of FASTQ files are essential to providing clean data to downstream analysis. By filtering raw reads with the trimmomatic (http://www.usadellab.org/cms/?page=trimmomatic), high-quality data were obtained. According to low quality sequence, these sequences were trimmed more than 30% of reads with quality less than Q20 and adapter (first 12bp of reads). After that, the clean reads were acquired, and the Homo sapiens hg19 genome reference and annotation GTF file was downloaded from the UCSC Genome Browser (https://genome.ucsc.edu/cgi-bin/hgGateway).

Using hisat2 (https://ccb.jhu.edu/software/hisat2/index.shtml) All clean reads were mapped to the Homo sapiens genome (mapping efficiencies (95%) for each paired end read). Then for the reading of transcripts, counting was performed with Htseq count (http://chipster.csc.fi/manual/htseq-count.html). Furthermore, using DESeq2 package (http://bioconductor.org/packages/release/bioc/html/DESeq.html) Differential expression analysis performed on three replicates of each normal and tumor ovarian samples.

### 2.3. Differential gene expression analysis based on the negative binomial distribution

#### 2.3.1. Checking the normalization of count

For evidence of systematic changes across experimental conditions, we applied suitable statistical approach to analysis and normalization of count data. In order to normalization of count data, we utilized DEseq, a method for differential analysis of count data, using shrinkage estimation for dispersions and fold changes to improve stability and interpretability of estimates. In order to better visualization, we transformed raw read counts using log2 transformation without quantile normalization Fig. 1 (Non-normalized log2 (counts+1) per sample) and then normalized raw read counts using the DESeq2 (log2 (normalized counts) per sample) Fig. 2.

**Fig. 1.**
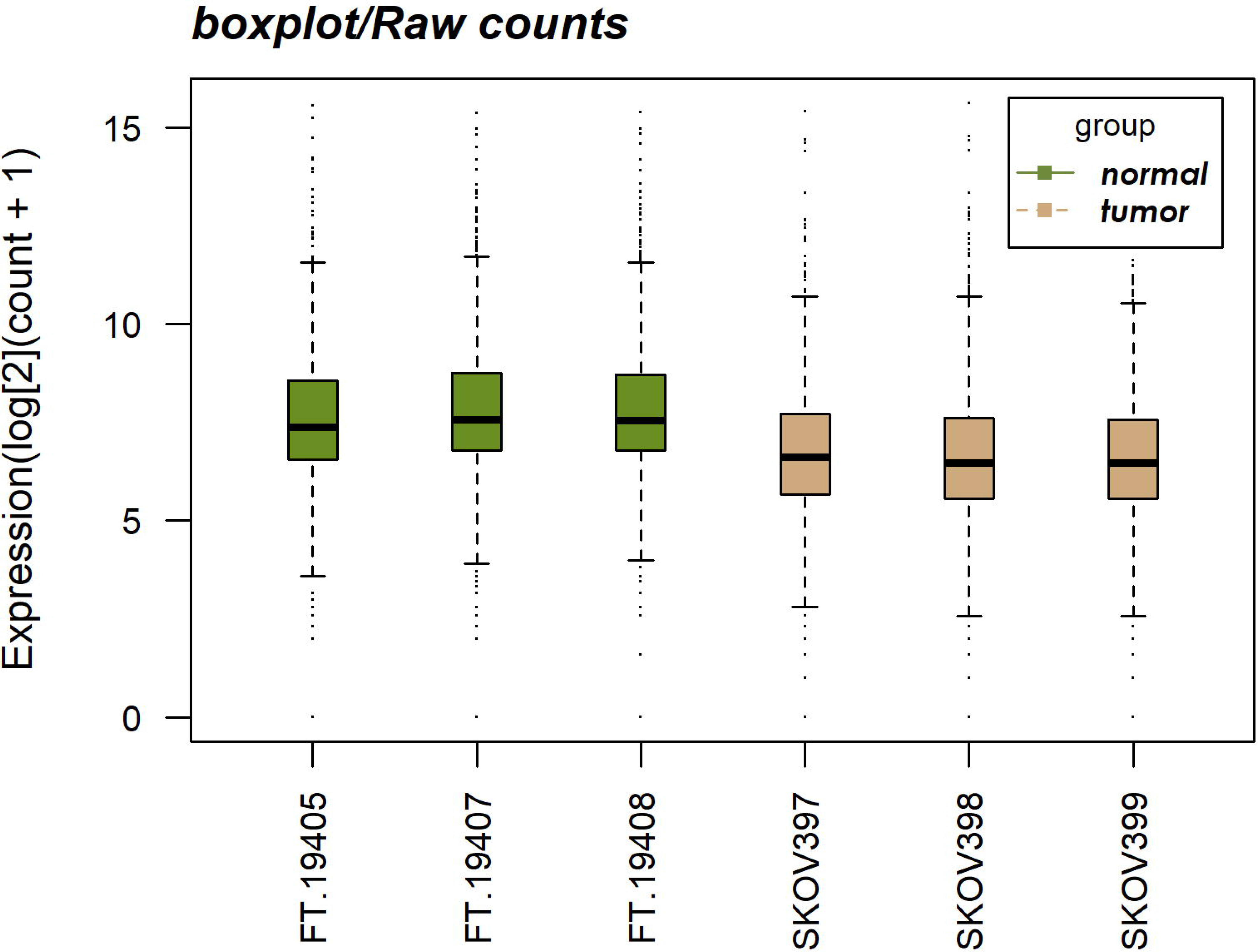
Box plots of non-normalized count (log2 (counts+1)) per sample.

**Fig. 2.**
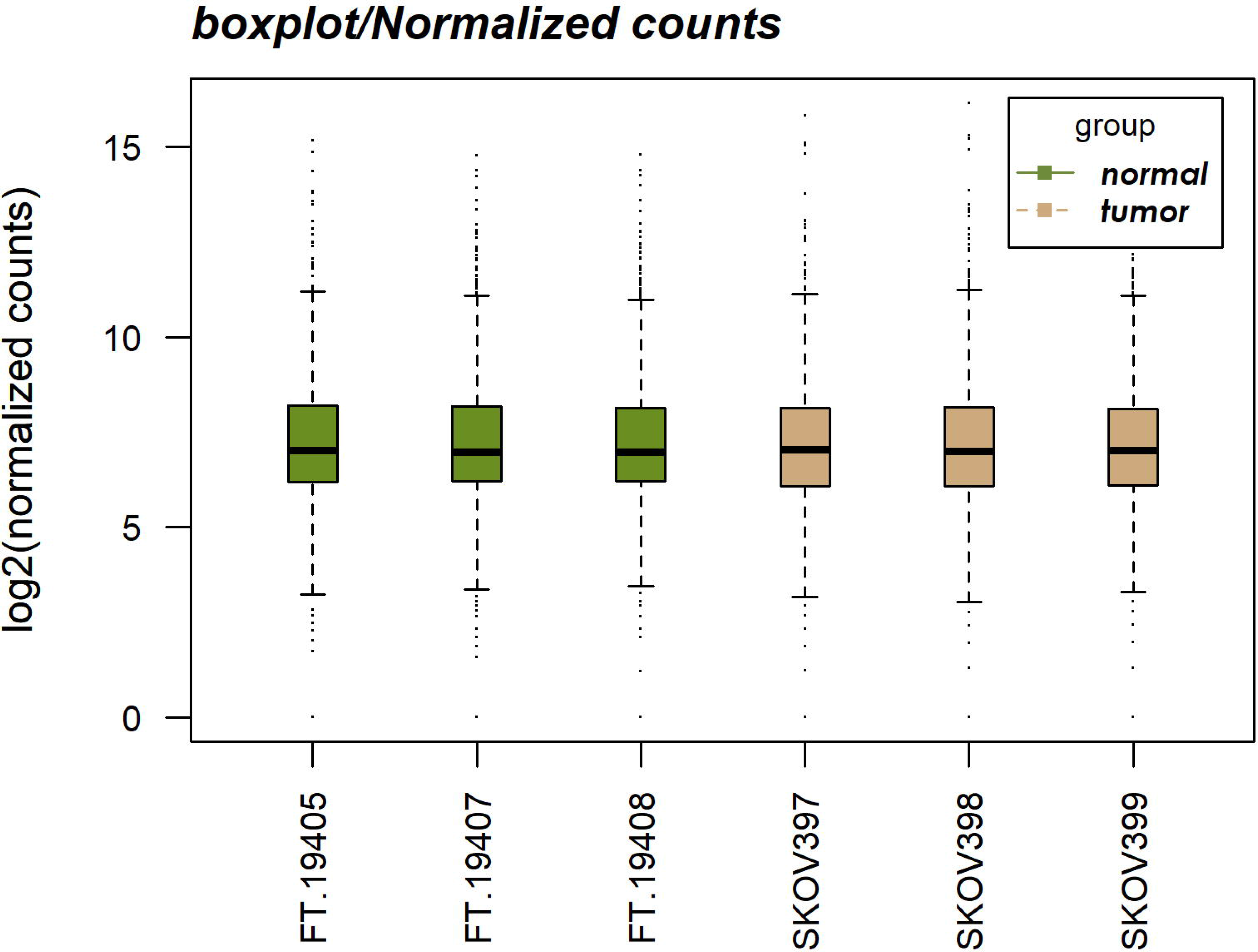
Box plots of normalized count (log2 (normalized counts)) per sample.

## 3. Results

### 3.1. Interpreting and visualization the DE analysis results

#### 3.1.1. Identifying the most Up-regulated and Down-regulated Genes

After normalization, DESeq2 was also used to screen the DEGs between ovarian tumor and normal groups. Totally, 1091 differential genes (padj<0.001) were obtained, 335 genes (padj<0.001, log2 FC >2) were significantly up-regulated, and 237 genes (padj<0.001, log2 FC <−2) were significantly down-regulated. The txt results were saved for subsequent analysis.

### 3.2. Hierarchical clustering of differentially expressed genes (DEGs)

To ensure that the selected genes are distinguished well between tumor and normal condition, we will perform a hierarchical clustering using the heatmap.2 function from the ggplot2 library. Rows correspond to genes, columns to samples. In the heatmap of Fig. 3, fifty differential genes based on the DESeq2 normalized gene expression with the lowest padj<0.001 were analyzed in the heatmap. The genes with similar expression patterns are clustered together. The up-regulated genes are in dark brown and the down-regulated genes are in golden.

**Fig. 3.**
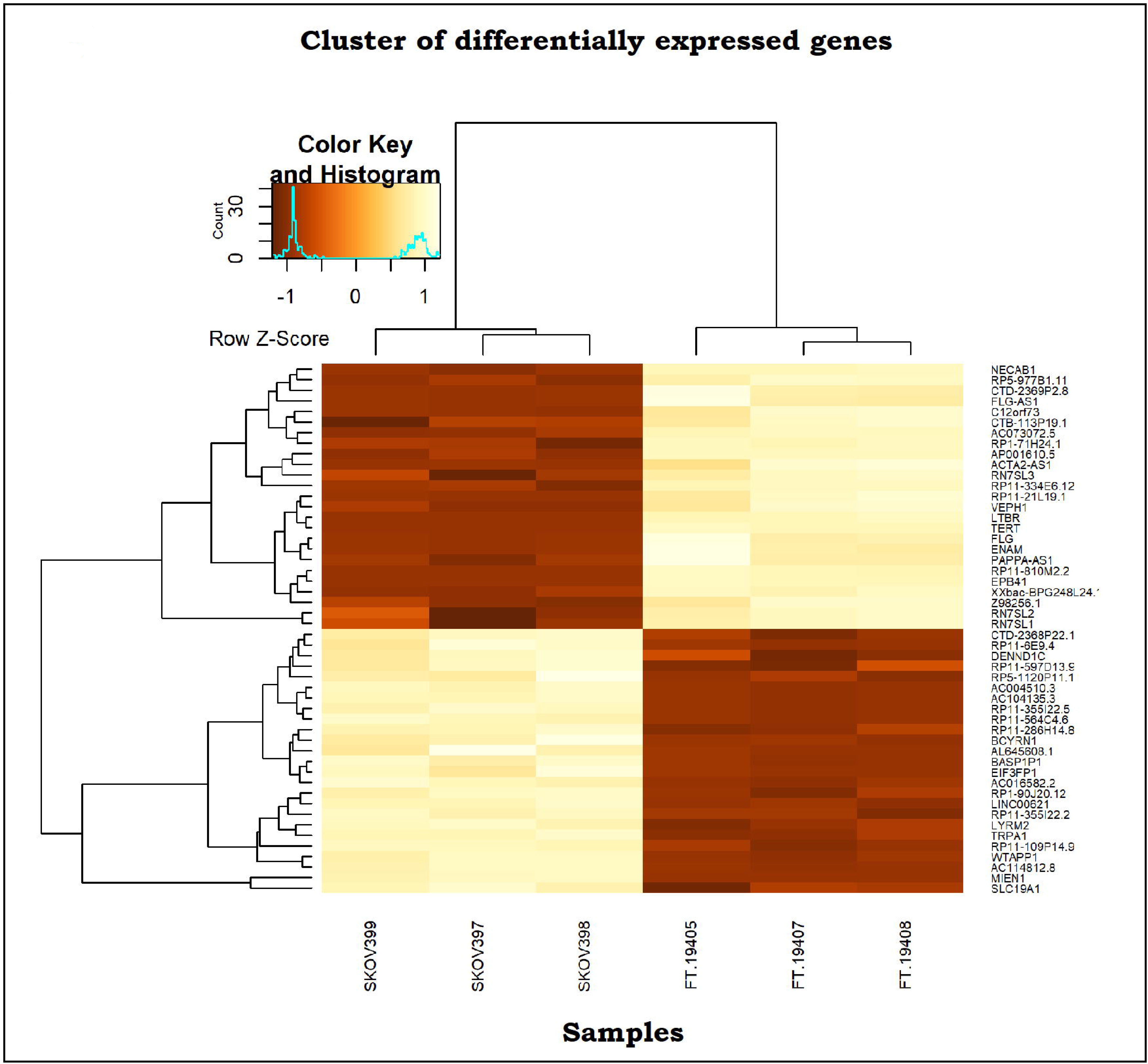
Heatmap across all the samples using the top 50 most DE genes between the normal and tumor ovarian groups.

### 3.3. MA-plot and Volcano plot

In DESeq2, the plotMA function shows the log2 FC attribution to given variable over the mean of normalized counts for an experiment with two-group comparison in the DESeqDataSet. The plot Fig. 4 represents each gene with a dot. The x-axis is the average expression over all samples, and the y-axis is the log2 Fold Change between normal and tumor group. Genes with an adjusted p value below a threshold (here 0.001) are shown in green. This plot demonstrates that only genes with a large average normalized count contain sufficient information to yield a significant call. The points, which fall out of the window, are plotted as open triangles pointing either up or down point..

**Fig. 4.**
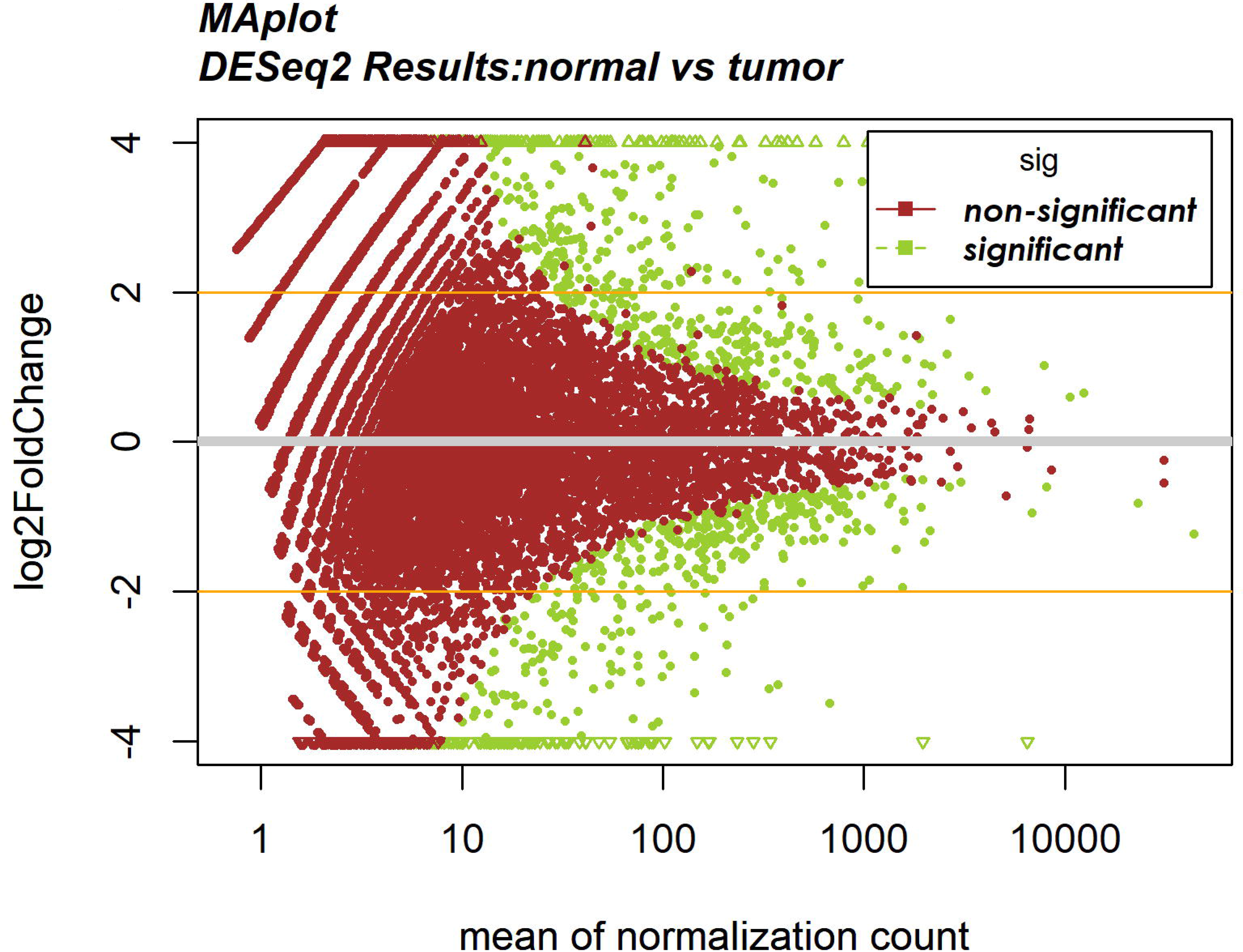
MA plot from Differential Expression genes between normal vs tumor ovarian groups. Oranges horizontal lines represents a log fold changes of −2 and 2.

In Fig. 5, 20 differential genes with the lowest adjusted p value were shown in the volcano plot. The y-axis corresponds to the mean expression value of negative log 10 (adjusted p-value), and the x-axis displays the log2 FC value. The yellow dots represent the up-regulated expressed genes (padj<0.001, log2 FC >2) between normal and tumor groups. The dark brown dots represent the genes whose expressions are down-regulated (padj<0.001, log2 FC < −2) between mentioned grupes.

**Fig. 5.**
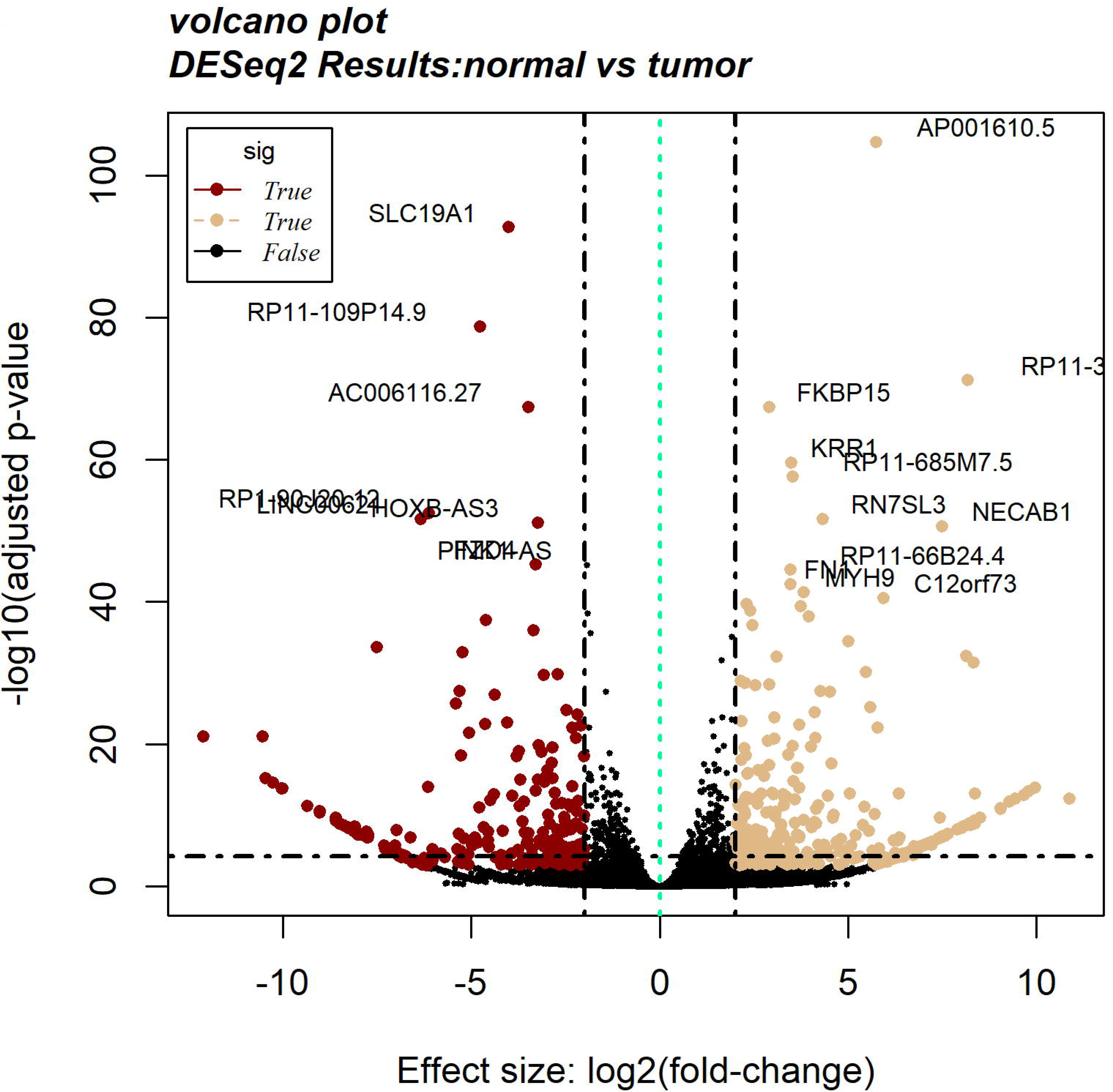
Volcano plot of the differentially expressed genes between normal vs tumor ovarian groups. Significantly down-regulated genes are in dark brown (sig=True), significantly up-regulated genes are in yellow (sig=True), non-significant genes are in black (sig=False). Black vertical lines highlight log fold changes of −2 and 2, while Black horizontal line represents a padj of 0.001.

### 3.4. Functional analysis of DEGs

To identify functional categories of differentially expressed genes, Gene Oncology (GO) enrichment analysis was performed using the R package, enrichR(https://cran.r-project.org/package=enrichR) provides an interface to the Enrichr database [39] hosted at (http://amp.pharm.mssm.edu/EnrichR). The groups with an padj<0.001 and gene counts more than two were examined. Enrichment of biological pathways supplied by enrichR as well.

### 3.5. Gene ontology (GO) enrichment analysis of differential expressed genes

Gene ontology (GO) was applied to identify characteristic biological attributes of RNA-seq rwa data. Separate Gene Ontology Enrichment Analysis using enrichR for up-regulated and down-regulated genes was performed. Furthermore, they were classified into different functional categories according to the GO term enrichment analysis for Biological process (BP) (Fig. 6), molecular function (MF) (Fig. 7) and cellular component (CC) (Fig. 8). 572 out of 1091 profiled DEGs assigned to 938 GO terms were enriched (padj <0.001). The top 12 significantly up-regulated and down-regulated GO categories are shown in (Table. 1 and Table. 2). These significantly enriched terms could help us a lot to further understand the role, which DEGs played in EOC.

**Fig. 6.**
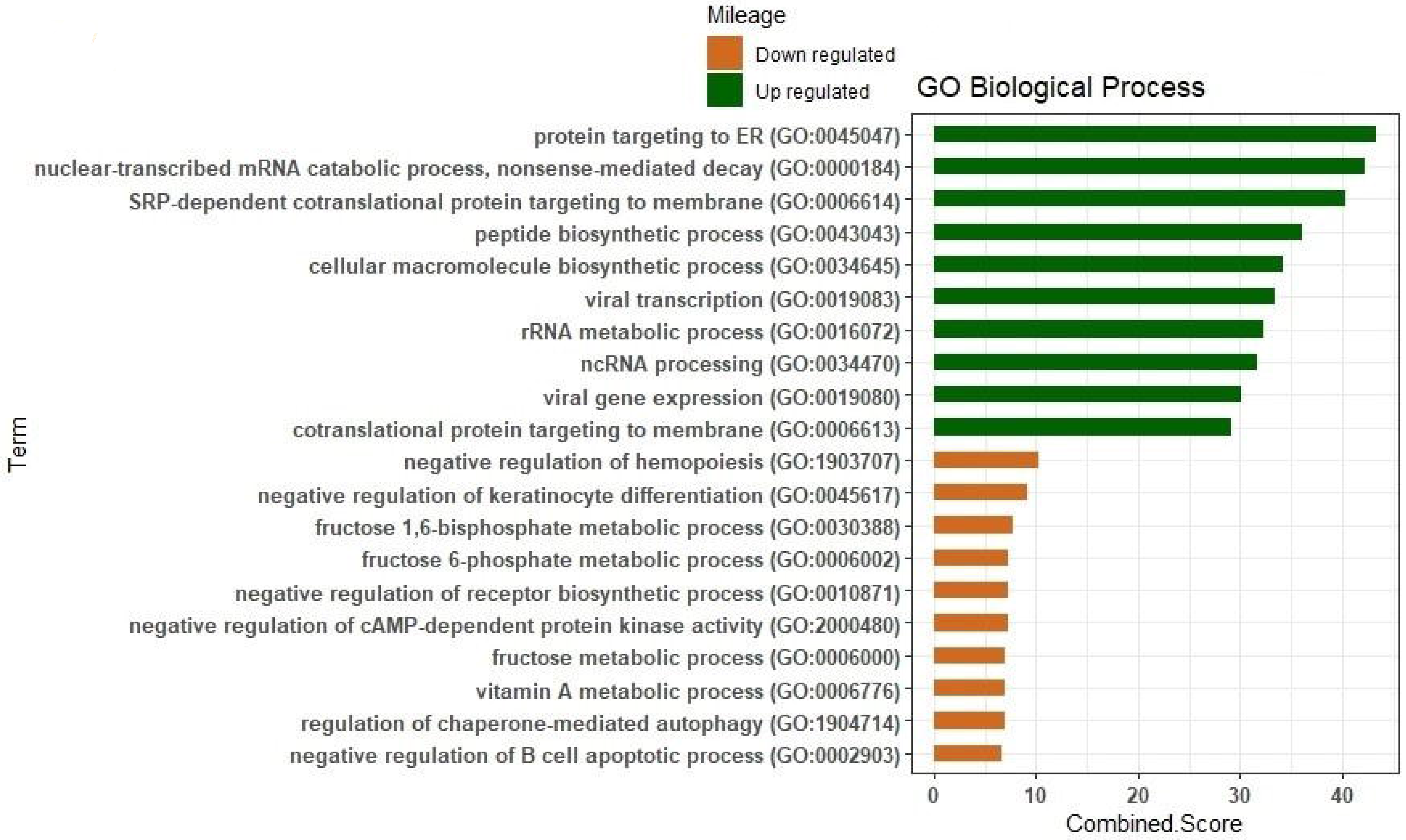
Gene Ontology enrichment analysis of biological processes for up- and down-regulated genes between normal vs tumor ovarian samples.

**Fig. 7.**
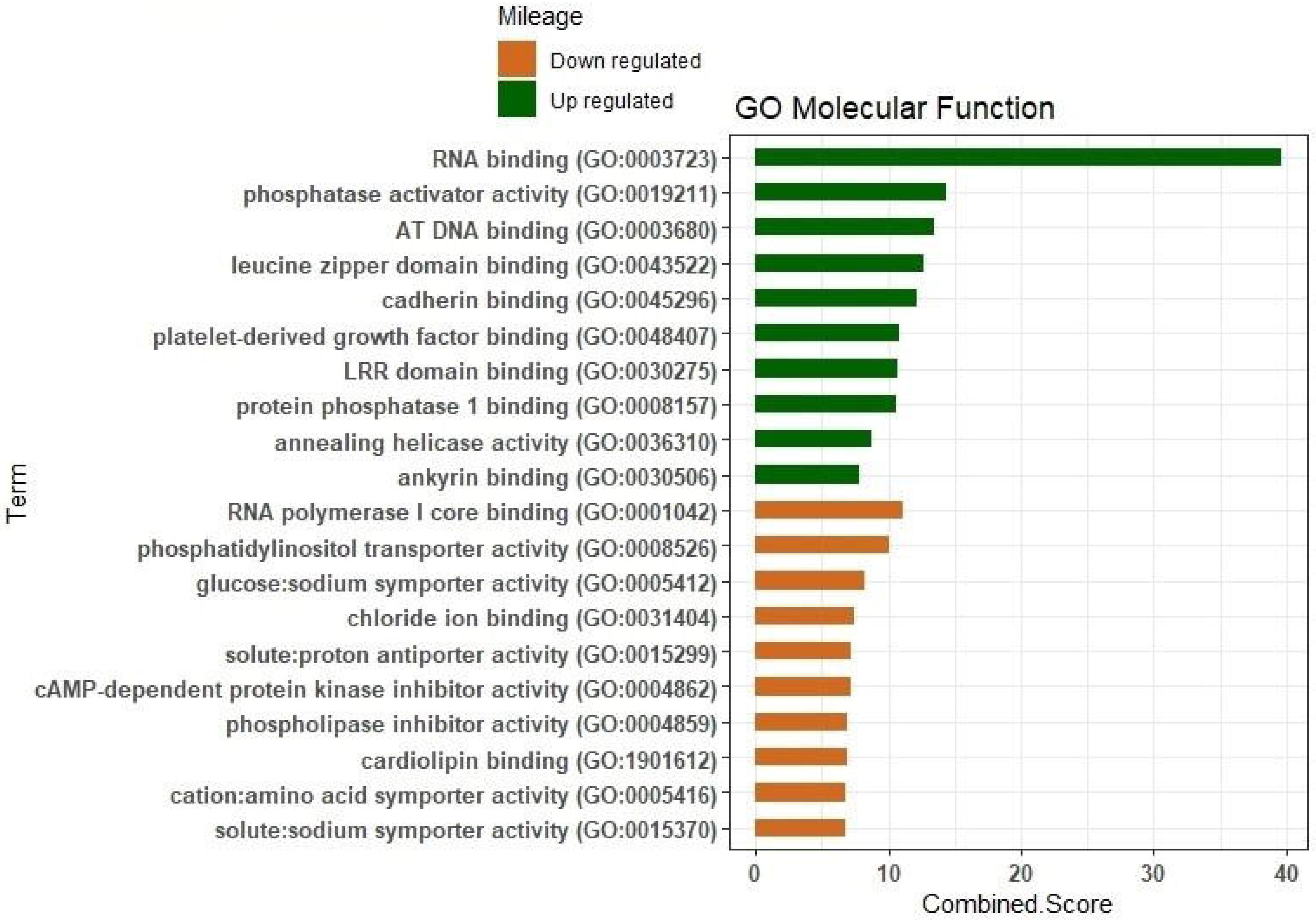
Gene Ontology enrichment analysis of molecular function for up- and down-regulated genes between normal vs tumor ovarian samples.

**Fig. 8.**
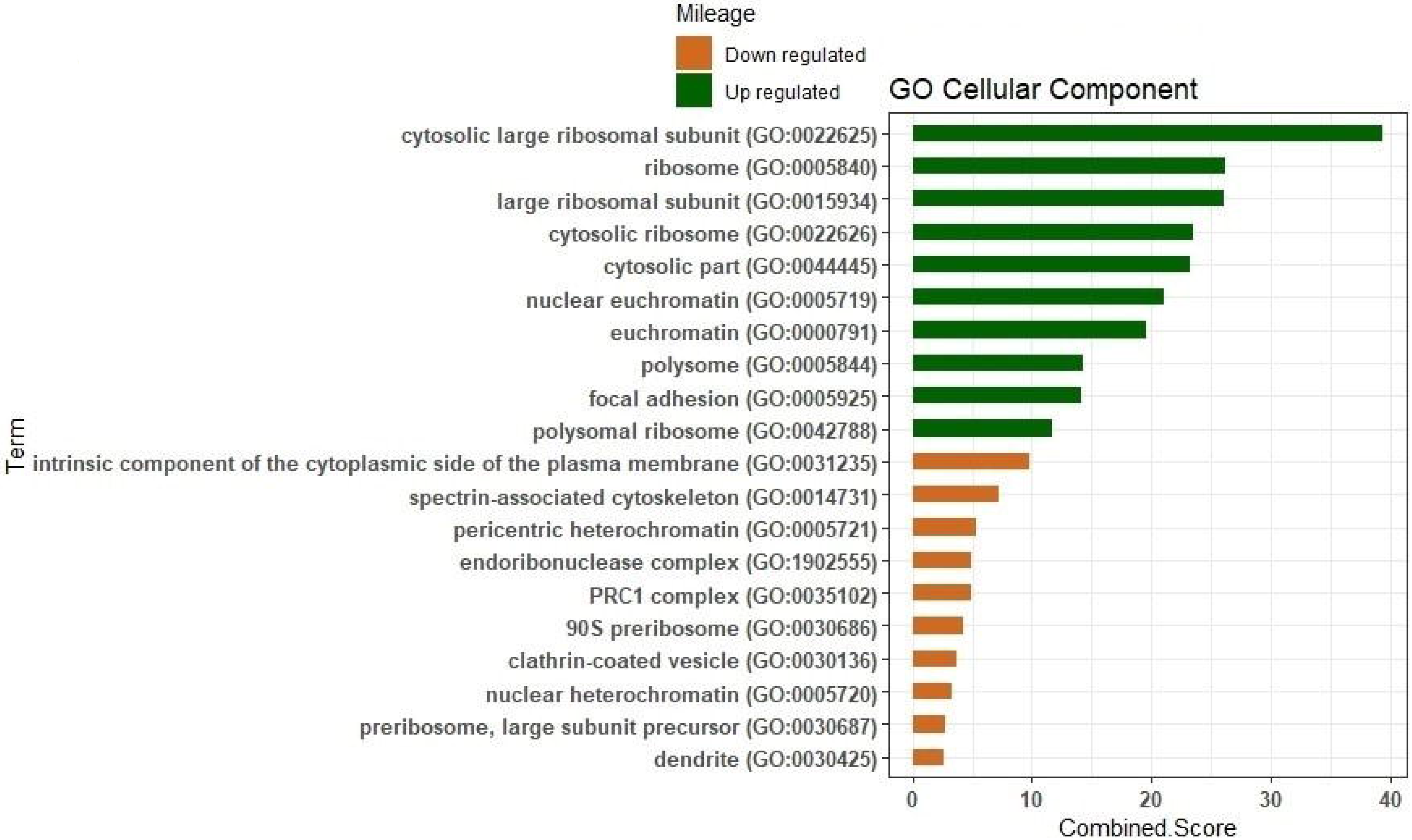
Gene Ontology enrichment analysis of cellular component for up- and down-regulated genes between normal vs tumor ovarian samples.

### 3.6. Kyoto Encyclopedia of Genes and Genomes (KEGG) pathway analysis

To investigate which DEGs is activated and suppressed in different class of pathways, gene expression information was mapped to the KEGG pathway. Pathway analysis and functional annotation for up- and down-regulated genes were performed using R program package enrichR. In total, 156 up-regulated genes (padj<0.001, log2 FC >2), and 82 down-regulated genes (padj<0.001, log2 FC < −2) were mapped to 238 KEGG pathways. The top 10 enriched pathways are shown in Fig. 9, and the results are shown in Table. 3. DEGs were highly clustered in several signaling pathways such as Ribosome, Focal adhesion, Glycolysis / Gluconeogenesis, Tryptophan metabolism, Apoptosis, Tryptophan metabolism, Pyruvate metabolism and Pyruvate metabolism.

**Fig. 9.**
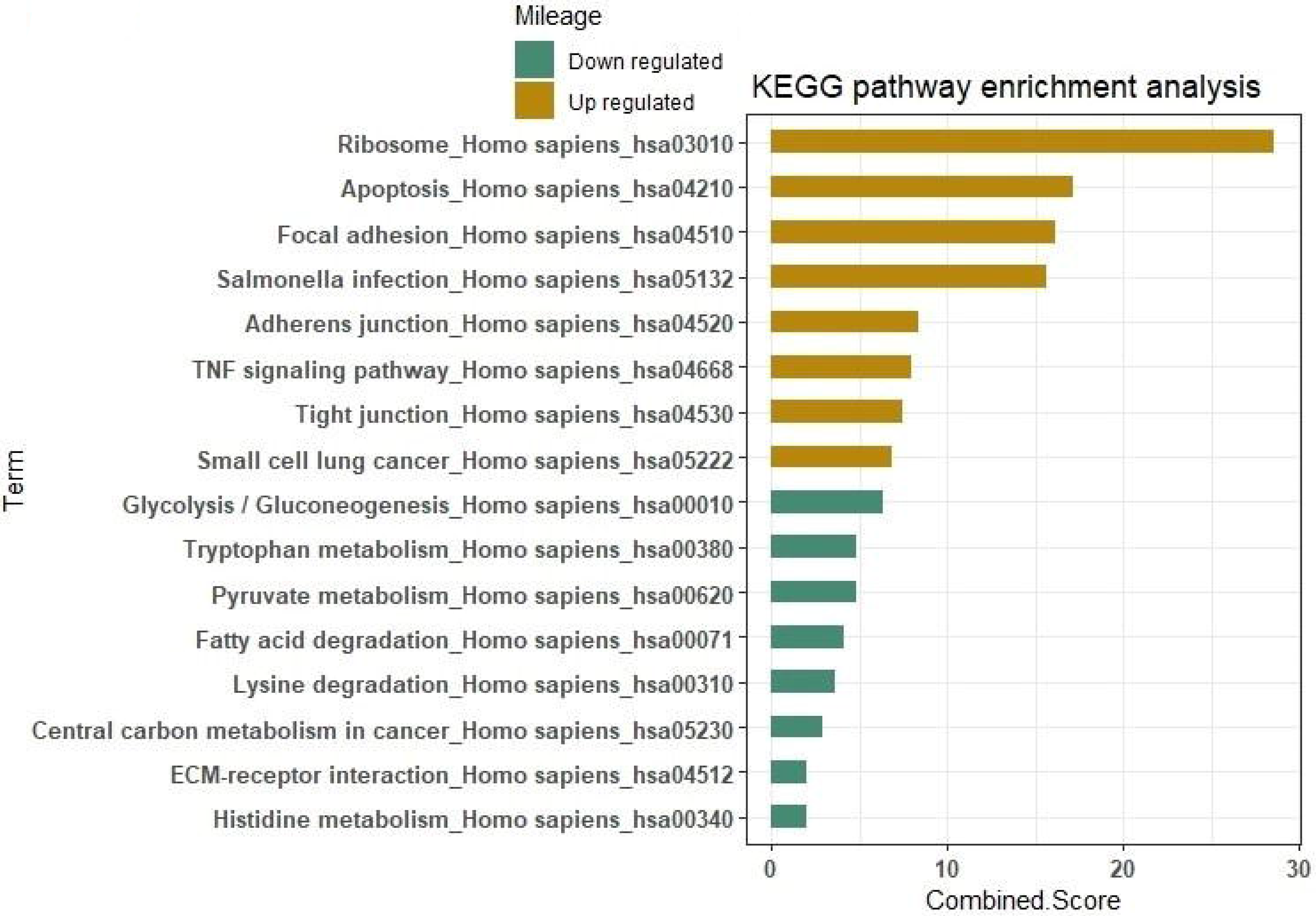
Kyoto Encyclopedia of Genes and Genomes (KEGG) pathway enrichment analysis for up- and down-regulated genes between normal vs tumor ovarian samples.

## 4. Discussion

Epithelial ovarian cancer (EOC) has the highest mortality rate among women's cancers due to the poor diagnosis, lack of the validated clinical applicable test to effective therapies [40]. Studies have shown that in order to achieve effective methods for early diagnosis and prevention of metastasis of ovarian cancer, it is important to study the molecular mechanisms of the carcinogenesis and development.

The purpose of current study is to investigate the novel genes involved in regulatory pathways in EOC using analyzing the RNA-Seq raw data and interpreting DEGs between normal and tumor ovarian samples, Ubuntu 17.10 (64-bit) and R programing environment (version 3.5.1) were used to extract DEGs information from Fastq data (SRR3289902). Many pivotal genes and pathways associated with ovarian cancer were identified in the present study. In the first step, 1091 DEGs (333 up-regulated and 273 down-regulated) were identified between normal and tumor samples. In the second step, 572 out of 1091 profiled DEGs (Padj<0.001) assigned and enriched to 938 GO terms, and then were classified into three groups cellular component (CC), biological process (BP) and molecular function (MF) using R program package (enrichR). In the third step, the up- and down-regulated DEGs were further clustered based on functions and signaling pathways with significant enrichment analysis.

The top 12 GO terms of the up- and down-regulated DEGs involved in BP, CC, and MF according to padj< 0.001 are shown in Table 1 and Table 2, respectively. The most significantly up-regulated DEGs were involved in the different GO terms such as protein targeting to ER, ontology; BP, RNA binding, ontology; MF, cytosolic large ribosomal subunit, ontology; CC (Fig. 6, Fig. 7, Fig. 8). The most significantly down-regulated DEGs were related to the GO terms negative regulation of hemopoiesis, ontology; BP, RNA polymerase I core binding, ontology; MF and intrinsic component of the cytoplasmic side of the plasma membrane, ontology; CC (Fig. 6, Fig. 7, Fig. 8).

In addition, KEGG pathway analysis demonstrated that the up-regulated DEGs were enriched in six significant pathways (Table 3 and Fig. 9), including Ribosome and Focal adhesion, whereas the down-regulated DEGs were enriched in six different pathways (Table. 3 and Fig. 9), including Glycolysis/Gluconeogenesis and Tryptophan metabolism. Among all GO terms, the most significantly up-regulated DEGs were RPL41, RBM25 and the most significantly down-regulated DEGs were HOXB8, IL17D, ERBB2, MIEN1 and SPTB. Furthermore, among all pathways, the most significantly up-regulated and down-regulated DEGs were RPL41, ALDH3A2, JUN, NFKBIA and TJP1.

In RNA polymerase I core binding term (GO: 0001042) Human epidermal growth factor receptor 2 (HER2, official name ERBB2) [41] was most up-regulated gene. This gene coded HER2 proteins, which is tyrosine kinase receptor in the epidermal growth factor (EGF) family and play a pivotal role in cell proliferation and tumor cell metastasis [42], [43], [44]. Overexpression of this gene has been reported in numerous cancers, including breast, ovarian, endometrial, salivary gland and gastric carcinomas [45], [46], [47]. HER2 are overexpressed in a significant percentage of epithelial ovarian cancers [48], [49]. In order to introduce this gene as biomarker, many studies have been made so that in many studies reported that HER2 expression was low in normal ovarian epithelium, while expressed highly in a variable percentage of epithelial ovarian cancer (11%-66%) [50]. In many study has been reported that the expression of HER2 may be a prognostic biomarker in non-serous ovarian cancer rather than serous for ovarian cancer [51].

Multiple endocrine neoplasia type 1 (MEN1) [52] was most down-regulated gene in intrinsic component of the cytoplasmic side of the plasma membrane term (GO: 0031235). This gene is an autosomal dominant cancer predisposition syndrome and encodes menin. It’s also a putative tumor suppressor associated with a syndrome known as multiple endocrine neoplasia type 1 [53], [54]. In several studies have been shown that menin either negatively regulates cell growth or participates in maintenance of genomic integrity [55], [56], [57], [58], [59].

In this study, the differentially expressed ribosomal Rps genes (RPL41, RPL3, RPL32, RPL13A, RPS25, RPS19, RPL14 and RPL36) were remarkable in ovarian cancer tumor. In protein targeting to ER (GO: 0045047) and among most of all GO term RPL41 ribosomal protein [60] was most up-regulated gene. This gene codes L4160S ribosomal protein, a small ribosomal peptide deregulated in tumors, is essential for mitosis and centrosome integrity and as a novel protein coding gene annotated in the GRCh38 human genome database [61]. It is possible that the lack of RPL41 in cells renders microtubule unstable and therefore premature centrosome splitting. Wang et al, [62] showed that the abnormal mitosis and disrupted centrosome associated with the RPL41 downregulation could be related to the malignant transformation

In leucine zipper domain binding (GO: 0043522), activating transcription factor 4 (ATF4) was upregulated. This gene is a major coordinator of tumor cell survival in stress and is commonly overexpressed in tumors [63]. Some investigations suggested that the ATF4 is a potential therapeutic target for cancer. There are many studies that show the close relationship between ATF4 and RPL41 [64]. Cells with RPL41 knockdown have significantly increased ATF4. RPL41 could also play a physiological role in regulating the cellular ATF4 level and induce rapid ATF4 degradation [65]. The RPL41 downregulation and deletions were mostly detected in human tumors. In the present study, RPL41 and ATF4 were up-regulated, maybe the augmentation of ATF4 can justify the increase of RPL41 in the most of GO terms in our study.

Most of up-regulated genes (59 gene) were found for RNA binding term (GO: 0003723). RBM25 that code RNA-binding protein was most up-regulated gene in this term and acted as a regulator of alternative pre-mRNA splicing, and it is involved in apoptotic cell death through the regulation of the apoptotic factor BCL2L1 isoform expression [66]. RBM25 is reported to promote apoptosis in HeLa cells by regulating the balance of pro- and anti-apoptotic transcripts of the BCL2L1 [67].

The Ribosome pathway and Glycolysis/Gluconeogenesis were identified as the most significantly enriched pathway in KEGG analysis. Fourteen of the most up-regulated genes (table.3, Fig. 9) were enriched in Ribosome pathway and three of the most down-regulated genes were enriched in Glycolysis / Gluconeogenesis pathway (table. 3, Fig. 9). ALDH3A2 [68] was most down-regulated gene in most KEGG pathways. This gene is a member of the aldehyde dehydrogenase (ALDH) gene family [69]. ALDH enzymes play a crucial role in epithelial homeostasis. Thus, deregulation of these enzymes is linked to multiple cancers such as breast, prostate, lung and colon cancers [70], [71], [72]. This enzyme breakdown fats and convert them into energy [73]. Chronic cell proliferation requires corresponding adjustment of energy metabolism in order to fuel rapid cell growth and division [74]. In addition, high glycolysis in tumor cells correlates with the degree of tumor malignancy, its ability to form metastasis and has been showed to be related to resistance against chemo- and radiotherapy treatments [75]. High levels of glucose are required to generate energy for the proliferation of cancer cells, Swa et al [76], showed that ALDH enzyme expression and activity may be associated with specific cell types in ovarian tumor tissues and vary according to cell states. Elucidating the function of the ALDH isozymes in lineage differentiation and pathogenesis may have significant implications for ovarian cancer pathophysiology [77], but in the present study, this gene has declined in most of the KEGG pathways. The reason for this controversy, which confirm these results, need to more experimental and computational studies. Our report may be of vital importance for investigating the complex interacting mechanisms underlying EOC carcinogenesis and designing specific treatments for patients with EOC.

According to World Health Organization (WHO) criteria, ovarian tumors can be classified as benign, low malignant potential or malignant. The histologic classification of ovarian carcinomas is based on morphologic criteria and corresponds to the different types of epithelial in the female reproductive system [78]. Unfortunately, the molecular mechanisms underlying ovarian carcinogenesis and histological differentiation remain unclear. However, current study provides a comprehensive bioinformatics analysis of the DEGs and pathways, which may be involved in designing and predicting of biomarkers associated with EOC.

## Conclusion

Final result of the present study, would promote an understanding of advanced EOC pathogenesis, and provide a number of valuable genes and pathways for further investigation of prognostic markers in advanced EOC. By using R software and Linux, we identified most significantly up- and down-regulated genes that could be the most related to the occurrence and development of EOC. Among them, 333 genes were up-regulated and 273 genes were down-regulated. Differential expression genes (DEGs) including RPL41, ALDH3A2, ERBB2, MIEN1, RBM25, ATF4, UPF2, DDIT3, HOXB8 and IL17D as well as the Ribosome pathway and Glycolysis/Gluconeogenesis pathway have had the potentiality to be used as targets for EOC diagnosis and treatment. The pivotal pathways and significant genes identified by us can be used for adaptation different EOC study. However, further molecular biological experiments, computational process and Gene co-expression network analysis are required to confirm the function of the identified genes associated with EOC. Because of the histologic classification of ovarian carcinomas, which is based on morphologic criteria and corresponds to the different types of epithelial in the female reproductive system, it is necessary to analyze and interpret individually each ovarian cancer dataset. In this case, significant genes and pathways for ovarian normal and tumor samples were identified, and urgent proceedings could be taken to identify the different stages of progression of EOC.

## Table captions

Table. 1. The top 12 enriched gene ontology terms of down-regulated DEGs involved in BP, CC, MF according to padj<0.001, log2 FC <− 2

Table. 2. The top 12 enriched gene ontology terms of up-regulated DEGs involved in BP, CC, MF according to padj<0.001, log2 FC >2

Table. 3. The top six enriched KEGG pathways (padj<0.001) of up-regulated and down-regulated differentially expressed genes associated with ovarian cancer.

## References

1. J. Krzystyniak, L. Ceppi, D.S. Dizon, M.J. Birrer. Epithelial ovarian cancer: the molecular genetics of epithelial. Ann. Oncol. 27 (2016), pp. i4–i10.

2. L.A Torre, B. Trabert, C.E. DeSantis, K.D. Miller, G. Samimi, et al. Ovarian cancer statistics. CA. Cancer J. Clin. 68 (2018), pp. 284–296.

3. A.D. Thor, R.H. Young, P.B. Clement. Pathology of the fallopian tube, broad ligament, peritoneum, and pelvic soft tissues. Hum. Pathol. 22 (1991), pp. 856–867.

4. V.M. Cho, E.K. Howell, Colvin. The Extracellular Matrix in Epithelial Ovarian Cancer—A Piece of a Puzzle. Front. Oncol. 5 (2015), pp. 245.

5. A.W. Kennedy, C.V. Biscotti, W.R. Hart, K.D. Webstar. Ovarian clear cell adenocarcinoma. Gynecol Oncol. 32 (1989), pp. 342–349.

6. U. Testa, E. Petrucci, L. Pasquini, G. Castelli, E. Pelosi. Heterogeneity and Progression, Clonal Evolution and Cancer Stem Cells. Medicines (Basel). 5 (2018), pp. Pii.E16.

7. E.L. Christie, D.D.L. Bowtell. Acquired chemotherapy resistance in ovarian cancer. Med. Oncol. (Northwood, London, England). 28 (2017), pp. viii13–viii15.

8. R. Archana, K.B. Simmons, C. Robert, J.r. Bast, The Emerging Role of HE4 in the Evaluation of Advanced Epithelial Ovarian and Endometrial Carcinomas. Oncology (Williston Park). 27 (2013), pp.548–556.

9. M. Denel-Bobrowska, M. Lukawska, I. Oszczapowicz, M. Marczak. Histological. Subtype of Ovarian Cancer as a Determinant of Sensitivity to Formamidine Derivatives of Doxorubicin in Vitro Comparative Studies with SKOV-3 and ES-2 Cancer Cell Lines. Asian. Pac. J. Cancer. Prev. 17 (2016), pp. 4223–4231.

10. S. Sarojini, A. Tamir, H. Lim, S. Li, S. Zhang. A. Goy, et al. Early Detection Biomarkers for Ovarian Cancer. J. Oncol. J Oncol. 2012 (2012), pp. 709049.

11. T.R. Adib, S. Henderson, C. Perrett, D. Hewitt D. Bourmpoulia, et al. Predicting biomarkers for ovarian cancer using gene-expression microarrays. Br. J. Cancer. 90 (2004), pp. 686–692.

12. R.L. Coleman, B.J. Monk, A.K. Sood, T.J. Herzog, et al. Latest research and clinical treatment of advanced-stage epithelial ovarian cancer. Nat. Rev. Clin. Oncol. 10 (2013), pp. 211–224.

13. Z. Wang, M. Gerstein, M. Snyder. RNA-seq: a revolutionary tool for transcriptomic. Nat. Rev. Genet. 10 (2009), pp. 57–63.

14. E. Kobayashi, Y. Ueda, S. Matsuzaki, T. Yokoyama, T. Kimura, et al. Biomarkers for Screening, Diagnosis, and Monitoring of Ovarian Cancer. J. Epidemiol. Biomarkers. Prev. 21 (2012), pp.1902–1912.

15. X. Wang, L. Han, L. Zhou, L. Wang, L.M. Zhang, Prediction of candidate RNA signatures for recurrent ovarian cancer prognosis by the construction of an integrated competing endogenous RNA network. J. Oncol. Rep. 40 (2018), pp. 2659–2673.

16. R. K. Kimberly, B.S. Montgomery. RNA Sequencing and Analysis. Cold. Spring. Harb. Protoc 11 (2015), pp. 951–69.

17. Y. Chu, D.R. Corey. RNA sequencing: platform selection, experimental design, and data interpretation. J. Nucleic. Acid. Ther. 22 (2012), pp. 271–274.

18. C.A. Maher, C. Kumar-Sinha, X. Cao, S. Kalyana-Sundaram, et al. Transcriptome sequencing to detect gene fusions in cancer. J. Nature. 458 (2009), pp. 97–101.

19. M.I. Love, W. Huber, S. Anders. Moderated estimation of fold change and dispersion for RNA-seq data with DESeq2. J. Genome. Biol. 15 (2014), pp. 550.

20. M.D Robinson, D.J. McCarthy, G.K. Smyth, edgeR: a Bioconductor package for differential expression analysis of digital gene expression data. Bioinformatics. 26 (2010), pp.139–140.

21. W. Huber, V.J. Carey, R Gentleman, S Anders, M Carlson, et al. Orchestrating high-throughput genomic analysis with Bioconductor. Nat. Methods. 12 (2015), pp.115–21.

22. D.J. McCarthy, Y. Chen, G.K. Smyth Differential expression analysis of multifactor RNA-Seq experiments with respect to biological variation. Nucleic. Acids. Res. 40 (2012), pp. 4288–4297.

23. T. Liu, Yu, N. Ding, F. Wang, S. Li, S. et al. verifying the markers of ovarian cancer using RNA⍰ seq data. Mol. Med. Rep. 12 (2015), pp.1125–30.

24. C.L. Barrett, C. DeBoever, K Jepsen, Saenz, C.C. Carson, D.A. Frazer, et al. systematic transcriptome analysis reveals tumor-specific isoforms for ovarian cancer diagnosis and therapy. Proc. Natl. Acad. Sci. U S A. 112 (2015), pp. E3050–E3057.

25. X. Yang, S. Zhu, L. Li, L. Zhang, S. Xian, et al. Identification of differentially expressed genes and signaling pathways in ovarian cancer by integrated bioinformatics analysis. Onco. Targets. Ther. 15 (2018), pp.1457–1474.

26. S. Anders, W. Huber. Differential expression analysis for sequence count data. Genome. biol. 11 (2010), pp. R106

27. J. Li, R. Tibshirani. Finding consistent patterns: A nonparametric approach for identifying differential expression in RNA-Seq data. Stat. Methods. Med. Res. 22 (2013), pp. 519–36.

28. M. Ashburner, C.A. Ball, J.A. Blake, D. Botstein, H. Butler, et al. Gene Ontology: tool for the unification of biology. Nat. Genet. 25 (2000), pp.25–9.

29. W. Huang da, B.T. Sherman, R.A. Lempicki. Bioinformatics enrichment tools: paths toward the comprehensive functional analysis of large gene lists. Nucleic. Acids. Res. 37 (2009), pp. 1–13.

30. M. Kanehisa S. Goto. KEGG: Kyoto Encyclopedia of Genes and Genomes. Nucleic Acids Res. 28 (2000), pp. 27–30.

31. X. wang, L. Han, L. Zhou, L. Wang, et al. Prediction of candidate RNA signatures for recurrent ovarian cancer prognosis by the construction of an integrated competing endogenous RNA network. Oncol. Rep. 40 (2018), pp. 2659–2673.

32. Y. Li, X. Xiao, X. Ji, B. Liu, C.I mos. RNA-seq analysis of lung adenocarcinomas reveals different gene expression profiles between smoking and nonsmoking patients. Tumour. Biol. 36 (2015), pp. 8993–9003.

33. W. Zhai, X.D. Yao, Y.F. Xu, B. Peng, H.M. Zhang, et al. Transcriptome profiling of prostate tumor and matched.normal samples by RNA-Seq. Eur. Rev. Med. Pharmacol. Sci. 18 (2014), pp.1354–60.

34. W. Yasui, N. Oue, R. Ito, K. Kuraoka, H. Nakayama, Search for new biomarkers of gastric cancer through serial analysis of gene expression and its clinical implications. J. Cancer. Sci. 95 (2004), p. 385–92.

35. J.M. Hao, J.Z. Chen, H.M. Sui, et al. A five-gene signature as a potential predictor of metastasis and survival in colorectal cancer. J. Pathol. 220 (2010), pp. 475–89.

36. W. Yasui, N. Oue, R. Ito, K. Kuraoka, H. Nakayama. Search for new biomarkers of gastric cancer through serial analysis of gene expression and its clinical implications. J. Cancer. Sci. 95 (2004), pp. 385–92.

37. S. Hyter, J. Hirst, H Pathak, Z.Y. Pessetto, D.C. Koestler, et al. Dev eloping a genetic signature to predict drug response in ovarian cancer. J. Oncotarget. 9 (2018), pp.14828–14848.

38. T. De Cristofaro, T. Di Palma, A. Soriano, A. Monticelli, O. Affinito, et al. Candidate genes and pathways downstream of PAX8 involved in ovarian high-grade serous carcinoma. J. Oncotarget. 7 (2016), pp. 41929–41947.

39. M.V. Kuleshov, M.R. Jones, A.D. Rouillard, N.F. Fernandez, Q. Duan, et al. Enrichr: a comprehensive gene set enrichment analysis web server. J. Nucleic. Acids. Res. 44 (2016), pp. W90–7.

40. R.L. Coleman, B.J. Monk, A.K. Sood, T.J. Herzog. Latest research and clinical treatment of advanced-tage epithelial ovarian cancer. Nat. Rev. Clin. Oncol. 10 (2013), pp. 211–224.

41. C. Gutierrez, R. Schiff. HER2: biology, detection, and clinical implications. Arch. Pathol. Lab. Med. 135 (2011), pp. 55–62.

42. W.G. McCluggage, N. Wilkinson. Metastatic neoplasms involving the ovary: a review with an emphasis on morphological and immunohistochemical features. J. Histopathology. 47 (2005), pp. 231–247.

43. Berchuck, A. Kamel, R. Whitaker, B. Kerns, G. Olt, et al. Overexpression of HER-2/neu is associated with poor survival in advanced epithelial ovarian cancer. J. Cancer. Res. 50 (1990), pp. 4087–4091.

44. H. Meden, D. Marx, T. Roegglen, A. Schauer, W. Kuhn, Overexpression of the oncogene c-erbB-2 (HER2/neu) and response to chemotherapy in patients with ovarian cancer. Int. J. Gynecol. Pathol. 17 (1998), pp. 61–65.

45. C.A. Bandera, H.W. Tsui, S.C. Mok, F.W. Tsui. Expression of cytokines and receptors in normal, immortalized, and malignant ovarian epithelial cell lines. J. Anticancer. Res. 23 (2003), pp. 3151–7.

46. F. Revillion, J. Bonneterre, J.P. Peyrat. ERBB2 oncogene in human breast cancer and its clinical significance. Eur. J. Cancer. 34 (1998), pp. 791–808.

47. D.J. Slamon, W. Godolphin, L.A. Jones, J.A. Holt, S.G. Wong, et al. Studies of the HER-2/neu proto-oncogene in human breast and ovarian cancer. J. Science. 244 (1989), pp. 707–12.

48. M.C. de Toledo, L.O. Sarian, L.F. Sallum, L.L. Andrade J Vassallo, et al. Analysis of the contribution of immunologically-detectable HER2, steroid receptors and of the "triple-negative" tumor status to disease-free and overall survival of women with epithelial ovarian cancer. Acta. Histochem. 116 (2014), pp. 440–7.

49. B.A. Goff, K Shy, B.E. Greer, H.G. Muntz, M Skelly, et al. Overexpression and relationships of HER-2/ neu, epidermal growth factor receptor, p53, Ki-67, and tumor necrosis factor alpha in epithelial ovarian cancer. Eur. J. Gynaecol. Oncol. 17 (1996), pp. 487–92.

50. M. Med, P. Sevelda, K. Czerwenka, K. Dobianer, H. Hanak, et al. DNA amplification of HER-2/neu and INT-2 oncogenes in epithelial ovarian cancer. J. Gynecol. Oncol. 59 (1995), pp. 321–326.

51. K.D. Steffensen, M.W. aldstrom, U. Jeppesen, E. Jakobsen, I. Brandslund, et al. The prognostic importance of cyclooxygenase 2 and HER2 expression in epithelial ovarian cancer. Int. J. Gynecol. Cancer. 17 (2007), pp. 798–807.

52. M.A. Gordon, H.M. Gundacker, J. Benedetti, J.S. Macdonald, J.C. Baranda, et al. Menin, the product of the MEN1 gene, is a nuclear protein. Proc. Natl. Acad. Sci. USA. 95 (1998), pp. 1630–4.

53. B. Valeria Busyginaa, E. Allen. Multiple Endocrine Neoplasia Type 1 (MEN1). Yale. J. Biol. Med. 79(2006), pp. 105–114.

54. L. Canaff, J.F. Vanbellinghen, H. Kaji, D. Goltzman, G.N. Hendy. Impaired transforming growth factor-TGF-) transcriptional activity and cell proliferation control of a menin in-frame deletion mutanβt associaβted with multiple endocrine neoplasia type 1 (MEN1). J. Biol. Chem. 287 (2012), pp. 8584–8597.

55. Y.S. Kim, A.L. Burns, P.K. Goldsmith, C Heppner, S.Y. Park, et al. Stable overexpression of MEN1 suppresses tumorigenicity of RAS. J. Oncogene. 18 (1999), pp. 5936–42.

56. Y. Yang, X. Hua. In search of tumor suppressing functions of menin. Mol. Cell. Endocrinal. 265-266 (2007), pp. 34–41.

57. S. Jin, H. Zhao, Y. Yi, Y. Nakata, A. Kalota, et al. Gewirtz, C-Myb binds MLL through menin in human leukemia cells and is an important driver of MLL-associated leukemogenesis. J. Clin. Invest. 120 (2010), pp. 593–606.

58. T. Wu, X. Zhang, X. Huang, Y. Yang, X. Hua. Regulation of cyclin B2 expression and cell cycle G2/m transition by menin. J. Biol. Chem. 285 (2010), pp.18291–300.

59. S.Y. Lin, S.J. Elledge, Multiple tumor suppressor pathways negatively regulate telomerase. Cell. 113 (2003), pp. 881–889.

60. “Entrez Gene: RPL41 ribosomal protein L41”

61. J. Klaudiny, H. von der Kammer, K.H. Scheit, Characterization by cDNA cloning of the mRNA of a highly basic human protein homologous to the yeast ribosomal protein YL41. J. Biochem. Biophys. Res. Commun. 187 (1992), pp. 901–6.

62. S. Wang, J. Huang, J. He, A. Wang, S. Xu, et al. RPL41, a small ribosomal peptide deregulated in tumors, is essential for mitosis and centrosome integrity. Neoplasia. 12 (2010), pp. 284–293.

63. K. Ameri, A.L. Harris, J. Int, Activating transcription factor 4 Biochem. Cell. Biol. 40(2008), pp.14–21.

64. H. Jiang, S. Hegde, B.L. Knolhoff, et al. targeting focal adhesion kinase renders pancreatic cancers responsive to checkpoint immunotherapy. Nat. Med. 22 (2016), pp. 851–60.

65. S. Wang, X. Xu, J. Zhang, D. He, Yan. Ribosomal protein RPL41 induces rapid degradation of ATF4, a transcription factor critical for tumour cell survival in stress. Ann. Oncol. 225 (2011), pp. 285–92.

66. Y. Tsujimoto. Role of Bcl-2 family proteins in apoptosis: apoptosomes or mitochondria. J. Genes. Cells. 3 (1998), pp. 697–707.

67. A.C. Zhou, A. Ou, E.J. Cho, S.C. Benz, Huang, et al. Novel splicing factor RBM25 modulates Bcl-x pre-mRNA 5' splice site selection. Mol. Cell. Biol. 28 (2008), pp. 5924–36.

68. C. Van den Hoogen, G. van der Horst, H Cheung, J.T. Buijs, R.C. Pelger, et al. The aldehyde dehydrogenase enzyme 7A1 is functionally involved in prostate cancer bone metastasis. Clin. Exp. Metastasis. 28 (2011), pp. 615–625.

69. P. Marcato, C.A. Dean, R. Araslanova, M. Gillis, M. Joshi, et al. Aldehyde dehydrogenase activity of breast cancer stem cells is primarily due to isoform ALDH1A3 and its expression is predictive of metastasis. J. Stem. Cells. 29 (2011), pp. 32–45.

70. P. Marcato, C.A. Dean, D. Pan, R Araslanova, M. Gillis, et al. Aldehyde dehydrogenase activity of breast cancer stem cells is primarily due to isoform ALDH1A3 and its expression is predictive of metastasis. Stem. Cells. 29 (2011), pp. 32–45.

71. Z.J. Liu, Y.J. Sun, J. Rose, Y.J. Chung, C.D. Hsiao, et al. Thfirst structure of an aldehyde dehydrogenase reveals novel interactions between NAD and the Rossmann fold. Nat. Struct. Biol. 4 (997), pp. 317–26.

72. S.A. Marchitti, D.J. Orlicky, C. Brocker, V. Vasiliou. Aldehyde dehydrogenase 3B1 (ALDH3B1): immunohistochemical tissue distribution and cellular-specific localization in normal and cancerous human tissues. J. Histochem. Cytochem. 58 (2010), pp. 765–783.

73. S.A. Marchitti, C. Brocker, D. Stagos, V. Vasiliou. Non-P450 aldehyde oxidizing enzymes: the aldehyde dehydrogenase superfamily. Expert. Opin. Drug. Metab. Toxicol. 4 (2008), pp. 697–720.

74. O. Warburg. On respiratory impairment in cancer cells. Science. 124 (1956), pp. 269–70.

75. R.H. Xu, H. Pelicano, Y. Zhou, J.S. Carew, L Feng, et al. Inhibition of glycolysis in cancer cells: a novel strategy to overcome drug resistance associated with mitochondrial respiratory defect and hypoxia. Cancer. res. 65 (2005), pp. 613–21.

76. B. Jackson, C. Brocker, D.C. Thompson, W. Black, K. Vasiliou, et al. Update on the aldehyde dehydrogenase gene (ALDH) superfamily. Hum. Genomics. 5 (2011), pp. 283–303.

77. Y.T. Saw, J. Yang, S.K. Ng, S. Liu, S. Singh, et al. Characterization of aldehyde dehydrogenase isozymes in ovarian cancer tissues and sphere cultures. BMC. Cancer. 1 (2012), pp. 329.

78. C. Meinhold-Heerlein, P.C. Fotopoulou, A. Harter, P. Kurzeder, A. Mustea, et al. The new WHO classification of ovarian, fallopian tube, and primary peritoneal cancer and its clinical implications. Arch. Gynecol. Obstet. 293 (2016), pp. 695–700.

