## Supplemental Table 1 for "Gene expression profiles and pathway enrichment analysis to identification of differentially expressed gene and signaling pathways in epithelial ovarian cancer based on high-throughput RNA-seq data"

| Term and GOID | DEGs | Adjusted p-value | Combined score | | Genes |
| --- | --- | --- | --- | --- | --- |
| negative regulation of hemopoiesis (GO:1903707) | BP | 0.9443 | | 10.32 | HOXB8;IL17D |
| negative regulation of keratinocyte differentiation (GO:0045617) | BP | 0.9443 | | 9.15 | CDSN |
| fructose 1,6-bisphosphate metabolic process (GO:0030388) | BP | 0.9443 | | 7.68 | FBP2 |
| fructose 6-phosphate metabolic process (GO:0006002) | BP | 0.9443 | | 7.33 |  |
| RNA polymerase I core binding (GO:0001042) | MF | 0.7527 | | 11.11 | ERBB2 |
| phosphatidylinositol transporter activity (GO:0008526) | MF | 0.7527 | | 10.01 | PITPNM3 |
| glucose:sodium symporter activity (GO:0005412) | MF | 0.7527 | | 8.26 | SLC5A9 |
| chloride ion binding (GO:0031404) | MF | 0.7527 | | 7.46 | NQO2 |
| intrinsic component of the cytoplasmic side of the plasma membrane (GO:0031235) | CC | 1 | | 9.82 | MIEN1;SPTB |
| spectrin-associated cytoskeleton (GO:0014731) | CC | 1 | | 7.234 | SPTB |
| pericentric heterochromatin (GO:0005721) | CC | 1 | | 5.30 | NCAPD3 |
| endoribonuclease complex (GO:1902555) | CC | 1 | | 4.93 | TSEN54 |

**Table. 2.**

| Term and GOID | DEGs | Adjusted p-value | Combined score | Genes |
| --- | --- | --- | --- | --- |
| protein targeting to ER (GO:0045047) | BP | 0.000000359 | 43.3031 | RPL41;RPL3;RPL32;RPL13A;RPS25;RPS19;RPL14;RPL36;RPL13;RPL26;RPL37;RPL29;RPS24;RPL19 |
| nuclear-transcribed mRNA catabolic process, nonsense-mediated decay (GO:0000184) | BP | 0.000000345 | 42.21 | UPF2;RPL41;RPL3;RPL32;RPL13A;RPS25;RPS19;RPL14;RPL36;RPL13;RPL26;RPL37;RPL29;RPS24;RP |
| SRP-dependent cotranslational protein targeting to membrane (GO:0006614) | BP | 0.000000345 | 40.38 | RPL41;RPL3;RPL32;RPL13A;RPS25;RPS19;RPL14;RPL36;RPL13;RPL26;RPL37;RPL29;RPS24;RPL19 |
| peptide biosynthetic process (GO:0043043) | BP | 0.00000108 | 36.03 | RPL41;EEF1A1P5;RPL3;WARS;RPL32;RPL13A;RPS25;RPS19;TNIP1;RPL14;RPL36;RPL13;RPL26;RPL3 |
| RNA binding (GO:0003723) | MF | 0.00000000009 | 39.57 | RBM25;RPL3;RPL32;HMGB2;PSIP1;YBX1;IFIT3;RPS19;HIST1H1D;RPL36;KIF1C;RPL37;HMGN2;HIST1H1B;HIST1H1C;CAST;DDX58;ACTN1;DNTTIP2;RPL13A;PPHLN1;GNL2;GTF2F1;RANGAP1;SMC1A;EEF1D;LUC7L3;MYH9;RPL26;SREK1;EZR;RPL29;PLEC;DHX8;SRRT;DDX21;PDCD11;TERT;PES1;UBC;RPL14;FLNA;RPL13;FLNB;SRSF11;RPL19;RBM39;JUN;PRPF38B;RPL41;KRR1;RCC2;PRRC2C;DEK;EEF2;RPS25;H1F0;MYBBP1A;ACO1;CALR;VIM;RPS24;WRAP53 |
| phosphatase activator activity (GO:0019211) | MF | 0.5918 | 14.35 | PPP1R15A;GTF2F1 |
| AT DNA binding (GO:0003680) | MF | 0.5476 | 13.411 | HAND2;HMGA1 |
| leucine zipper domain binding (GO:0043522) | MF | 0.5918 | 12.71 | DDIT3;ATF4 |
| cytosolic large ribosomal subunit (GO:0022625) | CC | 0.000001894 | 39.30 | RPL41;RPL3;RPL32;RPL14;RPL36;RPL13;RPL13A;RPL26;RPL37;RPL29;RPL19 |
| ribosome (GO:0005840) | CC | 0.0001502 | 26.14 | RPS25;RPL41;RPL32;RPS19;RPL36;RPL13;RPL13A;RPL26;RPL19 |
| large ribosomal subunit (GO:0015934) | CC | 0.000001894 | 26.10 | RPL41;RPL3;RPL32;RPL14;RPL36;RPL13;RPL13A;RPL |
| cytosolic ribosome (GO:0022626) | CC | 0.000001894 | 23.40 | RPL41;RPL3;RPL32;RPL13A;RPS25;RPS19;RPL14;RPL |

**Table. 3.**

| Pathway and ID | DEGs | Adjusted p-value | Combined score | Genes |
| --- | --- | --- | --- | --- |
| Ribosome_Homo sapiens_hsa03010 | up | 0.00001206 | 28.59 | RPL41;RPL3;R  PL32;RPL13A;RPS25;RPS19;RPL14;RPL36;RPL13;RPL26;RPL37;RPL29;RPS24;RPL19 |
| Focal adhesion_Homo sapiens_hsa04510 | up | 0.008411 | 16.34 | JUN;COL4A1;ACTN1;COL6A2;COL6A1;FN1;FLNA;FLNB;FLNC;ACTB;BIRC3;ACTG1 |
| Salmonella infection_Homo sapiens_hsa05132 | up | 0.007583 | 15.66 | TJP1;JUN;MYH9;FLNA;FLNB;FLNC;ACTB;ACTG1 |
| Apoptosis_Homo sapiens_hsa04210 | up | 0.02333 | 13.98 | NFKBIA;JUN;GADD45A;DDIT3;LMNA;SEPT4;ACTB;ATF4;BIRC3;ACTG1 |
| Adherens junction_Homo sapiens_hsa04520 | up | 0.2350 | 8.38 | TJP1;ACTN1;MLLT4;ACTB;ACTG1 |
| Glycolysis / Gluconeogenesis_Homo sapiens_hsa00010 | down | 0.9572 | 6.32 | ALDH3A2;LDHA;FBP2 |
| Tryptophan metabolism_Homo sapiens_hsa00380 | down | 0.9572 | 4.79 | ALDH3A2;GCDH |
| Pyruvate metabolism_Homo sapiens_hsa00620 | down | 0.9572 | 4.78 | ALDH3A2;LDHA |
| Fatty acid degradation_Homo sapiens_hsa00071 | down | 0.9572 | 4.14 | ALDH3A2;GCDH |
| Lysine degradation_Homo sapiens_hsa00310 | down | 0.9572 | 3.66 | ALDH3A2;LDHA |
